# Multiple sounds degrade the frequency representation in monkey inferior colliculus

**DOI:** 10.1101/2020.07.03.187021

**Authors:** Shawn M. Willett, Jennifer M. Groh

## Abstract

How we distinguish multiple simultaneous stimuli is uncertain, particularly given that such stimuli sometimes recruit largely overlapping populations of neurons. One commonly proposed hypothesis is that the sharpness of tuning curves might change to limit the number of stimuli driving any given neuron when multiple stimuli are present. To test this hypothesis, we recorded the activity of neurons in the inferior colliculus while monkeys made saccades to either one or two simultaneous sounds differing in frequency and spatial location. Although monkeys easily distinguished simultaneous sounds (∼90% correct performance), the frequency selectivity of inferior colliculus neurons on dual sound trials did not improve in any obvious way. Frequency selectivity was degraded on dual sound trials compared to single sound trials: neural response functions broadened, and frequency accounted for less of the variance in firing rate. These changes in neural firing led a maximum-likelihood decoder to perform worse on dual sound trials than on single sound trials. These results fail to support the hypothesis that changes in frequency response functions serve to reduce the overlap in the representation of simultaneous sounds. Instead, these results suggest that alternative possibilities, such as recent evidence of alternations in firing rate between the rates corresponding to each of the two stimuli, offer a more promising approach.

**Graphic Abstract:** How sensory representations encode multiple stimuli despite coarse coding is unknown. Using a maximum likelihood decoder operating on the spike count response patterns of monkey inferior colliculus neurons, we show a marked reduction in decoding accuracy when two sounds are presented compared to one. The decoding was inferior to the behavioral performance of the animals, and thus suggests the presence of alternative coding strategies.

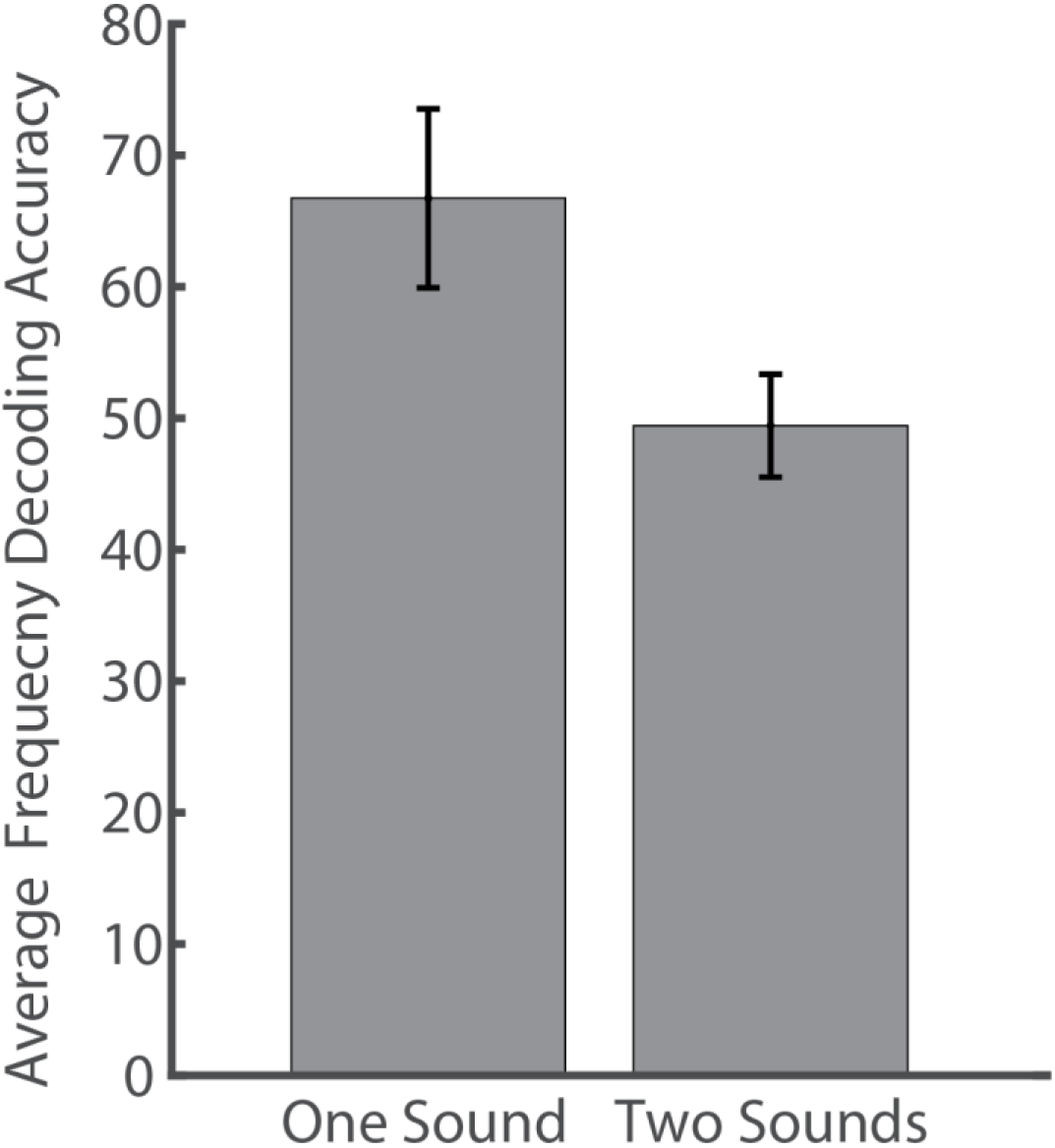

## Introduction

Natural environments contain numerous stimuli. How the brain simultaneously encodes multiple stimuli is not well understood. A key problem is that when more than one stimulus falls in a given neuron’s receptive field, it is not clear how the firing rate of that neuron can simultaneously encode both items. Two possible (and potentially complementary) solutions to this multiplicity conundrum have been proposed: 1) that receptive fields effectively shrink or shift to limit the number of stimuli within them, leading to more disparate populations representing each stimulus, and/or 2) that fluctuating activity patterns allow individual neurons to contribute to encoding only one stimulus at any given time. Recently Caruso *et al.* (2018) discovered neurons in the monkey inferior colliculus (IC) that display alternations in firing rates between those evoked by individual stimuli, supporting the fluctuating activity hypothesis (Caruso *et al.*, 2018). However, whether the underlying frequency response functions of these neurons may also shrink or shift (reducing the overlap in the potentially responsive population) remains unknown. Here, we evaluated whether and how the presence of two sounds might alter the frequency response properties of neurons in the IC.

The relevant physical cues to consider are several. The grouping or segregation of sound wave(s) into the percept of one vs. multiple sound sources depends on the timing, location, and frequency content of those sound wave(s), a phenomenon known as auditory scene analysis (Bregman, 1990). In most cases, at least two of these three factors must differ for sound sources to be perceived as distinct. When sound waves are simultaneous, they must therefore differ in both frequency and location to be perceived as arising from distinct underlying events (Blauert, 1983). The underlying neural apparatus responsible for such perceptual segregation must therefore have the ability to preserve information about both sounds on the basis of the combination of location and frequency content.

The monkey IC is an ideal structure to probe how neurons code concurrent stimuli differing in frequency content and location. Nearly every auditory signal converges to the IC prior to ascending to the thalamus (Aitkin & Phillips, 1984), so information must be preserved at this stage in order to be available to higher areas (for review see Winer & Schreiner, 2005). Causal studies have confirmed the IC is critical in both sound localization and frequency discrimination (Jenkins & Masterton, 1982; Kelly & Kavanagh, 1994; Pages *et al.*, 2016).

However, the representations of sound frequency and location in the monkey IC appear too coarse to account for monkeys’ perceptual abilities in any obvious way. Monkeys can distinguish a 2000 Hz tone from another tone in the 2020-2060 Hz range (Sinnott *et al.*, 1987). We have rarely observed or seen reports of individual primate IC neurons for which such a small 20-60 Hz (0.03 octave) change in frequency would cause it to stop (or begin) responding or significantly change its firing rate (Groh *et al.*, 2003; Bulkin & Groh, 2011),. Instead, most neurons in the primate and cat IC appear to respond over a range measured in octaves at above-threshold sound intensities (Ryan & Miller, 1978; Calford *et al.*, 1983; Ramachandran *et al.*, 1999; Zwiers *et al.*, 2004; Versnel *et al.*, 2009); see also Joris *et al.* (2011). At the population level, a given pure tone can activate anywhere from 20-80% of frequency modulated multi-unit recording sites in the monkey (Bulkin & Groh, 2011).

Sound localization behavior is also more precise than the neural code would seem to permit if governed by receptive field size. Both humans and monkeys can localize individual sounds with saccades to an accuracy of ∼4-6 degrees in the horizontal dimension (Jay & Sparks, 1990; Metzger *et al.*, 2004), but neurons tend not to have circumscribed spatial receptive fields at all. Instead, they respond monotonically as a function of increasing sound eccentricity over a broad range of space (a hemifield or more, Groh *et al.*, 2003; Zwiers *et al.*, 2004), as seen in other mammalian species (McAlpine & Grothe, 2003; Grothe *et al.*, 2010).

The coarseness of this coding is not necessarily a problem if only one sound is present at a time. In such cases, so long as each stimulus evokes activity in a slightly different population of neurons, simple read out mechanisms are potentially able to identify the stimulus with high resolution on the basis of the profile of activity across the population (Eurich & Schwegler, 1997). Indeed, the perceptual difference limens in the literature stem from studies in which sounds are presented individually. Coarse coding is also well known in motor systems, where a large proportion of the neural population may contribute to coding a given movement (Georgopoulos *et al.*, 1986; Lee *et al.*, 1988). However, only one movement will happen at any one time, whereas sensory systems are often charged with preserving information about multiple stimuli. Two concurrent sounds can be detected and/or localized by humans (Perrott, 1984a; b; Best *et al.*, 2004; Zhong & Yost, 2017) and monkeys (Caruso *et al.*, 2018), implying that the IC neural population and other auditory structures must somehow overcome these coding constraints.

Could the answer lie in some form of reduction in the coarseness of coding? The vibration of the basilar membrane elicited by one (monaural) tone frequency is altered by the inclusion of a second tone (two tone suppression, e.g. Ruggero *et al.*, 1992). Central response functions are not immutable either (Weinberger, 1995; Eggermont, 2011; David, 2018) – similar suppression of responses has also been observed in auditory cortex for either simultaneous or sequential stimuli (Schwarz & Tomlinson, 1990; Brosch & Schreiner, 1997) Computational analyses of responses to harmonic tone complexes further indicate the presence of inhibitory modulation of response patterns (Fishman *et al.*, 2013; Su & Delgutte, 2020), and auditory attention can alter auditory cortical frequency response patterns as well (Fritz *et al.*, 2007; Connell *et al.*, 2014). Spatial response functions can also be modulated. Auditory cortical units show broad selectivity in cats under anesthesia (Middlebrooks & Pettigrew, 1981; Imig *et al.*, 1990; Middlebrooks *et al.*, 1994) but become more selective when the cat is awake and/or performing a spatial auditory task (Mickey & Middlebrooks, 2003; Lee & Middlebrooks, 2011). Indeed, it is also well known that anesthesia affects the temporal response profile in marmoset monkey (Wang *et al.*, 2005; Bendor & Wang, 2007). These findings have parallels in other sensory systems, such as surround suppression in vision (Hartline & Ratliff, 1957; Chettih & Harvey, 2019) and changes to visual cortical object selectivity associated with experience (Freedman *et al.*, 2005).

These studies provide proof of principle that the selectiveness of neural response functions *could* change but do not show whether any such potential changes in selectivity are adequate to account for perceptual performance with multiple sounds. Indeed, previous work in the anesthetized rabbit IC found the presence of a concurrent sound distorted the shape of spatial response functions and led to a *decrease* in neural sensitivity to target position (Day *et al.*, 2012). However, changes in frequency response functions could also contribute to preserving information about multiple sounds (Blauert, 1983). The present study therefore investigates how frequency response functions in the monkey IC are modulated by the presence of an additional sound differing in frequency and location, and evaluates whether any such changes are beneficial to decoding stimulus information from the population.

Single neuron activity was recorded from the IC while monkeys made saccades to the positions of one or two simultaneously presented sounds differing in frequency by 0.25-2 octaves and in location by 24 degrees. Monkeys performed at least 86% correct on Dual sound trials for all frequency separations. Frequency response functions were often altered when a second sound was presented but did not appear to shrink or shift. Rather, most neurons actually broadened their frequency response functions. A maximum-likelihood decoder used to decode the stimulus frequency on a held-out trial performed slightly worse when the held-out trial was a Dual sound trial than a Single sound trial. Thus, the frequency selectivity modulations due to the presence of an additional sound seem to limit the available stimulus frequency information encoded by the population of IC neurons.

We conclude that changes to the frequency response properties of IC neurons in the presence of multiple sounds are unlikely to solve the multiplicity problem. This finding leaves standing an interesting alternative hypothesis: that coding remains coarse but that fluctuating activity patterns allow individual neurons to contribute to encoding only one stimulus at any given time (Fitzpatrick *et al.*, 1997; Caruso *et al.*, 2018). Future work to investigate how this alternative form of multiplicity coding may operate is likely to prove a more fruitful avenue to advancing our understanding of how the brain preserves information about multiple sounds.

## Methods

### General

All procedures were approved by the Duke University Institutional Animal Care and Use Committee. Two adult, female, rhesus monkeys (Monkey Y – 7 years old at start of recordings; Monkey N – 12 years old at start of recordings) participated in this study. General procedures have been previously described in greater detail (Groh *et al.*, 2001; Caruso *et al.*, 2018). Briefly, under aseptic conditions a head post and a scleral search coil were surgically implanted (Robinson, 1963; Judge *et al.*, 1980). After recovery monkeys were trained to perform a saccades-to-sound task as described below (Jay & Sparks, 1990; Metzger *et al.*, 2004; Caruso *et al.*, 2018). Once the task was learned, a recording chamber (Crist Instruments) was implanted over the IC based on stereotaxic coordinates and was confirmed after implantation with structural MRI (Groh *et al.*, 2001). The angle of the chamber allowed for access to both the right and left IC; all but one neuron were recorded from the right IC.

### Sound stimuli

All sounds consisted of bandpass filtered noise. Sounds were frozen for each experimental session; that is, for any given experimental session every instance of the sound stimulus of a given frequency involved the same waveform. The center frequencies were equally spaced in 0.25 octave intervals: 420, 500, 595, 707, 841, 1000, 1189, 1414, 1682 and 2000 Hz, and a proportional bandwidth of +/− 10% of the center frequency (−0.15 to +0.13 octaves). All sounds were initiated with a 10 ms on ramp and were generated at a sampling rate of 11 kHz. These sounds were chosen due to the large proportion of IC neurons responsive at these frequencies (Bulkin & Groh, 2011), increasing the chance that more than one sound would significantly drive any recorded neuron.

All sounds were generated using proprietary software (Beethoven, RYKLIN Software Inc.; SoundMax Integrated Audio HD sound card) and passed through appropriate amplifiers (Tucker Davis Technologies, SA1 Stereo Amplifier) to two speakers (Bose Acoustimass Cube Speakers) matched in their frequency response characteristics. Speakers were positioned at −12° and +12° in the horizontal azimuth and at 0° in elevation at a distance approximately 1 meter from the monkey. Sounds were calibrated using a microphone (Bruel and Kjaer 2237 sound level meter) at the position where the monkey’s head would be during the experiment. Sounds were calibrated to be 50 dB SPL when presented in isolation; these stimuli resulted in approximately 53-55 dB SPL sound reaching the monkey’s head on Dual sound trials. The intensity difference is not expected to have substantive effects on neural activity or behavior (Caruso *et al.*, 2018). We kept the sound source signals constant across conditions because in the real world simultaneously occurring sounds are louder than when the component sounds are presented in isolation. All experiments were conducted in an anechoic chamber; the wall and ceilings were lined with echo attenuating foam (Sonex) and the floor was carpeted.

### Task Conditions

#### Task overview

Monkeys performed three different trial types (Figure 1A & B): 1) Single sound trials involving one bandpass noise stimulus at a given center frequency and location; 2) Dual sound trials involving two bandpass noise stimuli differing in both frequency and location; and 3) Control trials in which two bandpass noise frequencies were presented at the same location. These control trials were spectrally equivalent to Dual sound trials, but spatially they were single target trials. For all three trial types, monkeys were rewarded for making saccades to the locations involved, i.e. two sequential saccades on the Dual sound trials and a single saccade on the single sound and control trials. Monkey Y was extensively trained on the single sound and Dual sound trials before the control trials were introduced, and performed this control task correctly on the first day, indicating that the monkey understood the importance of making a saccade to the sound source location on the correct side rather than simply responding with two saccades to memorized locations for trials with two center frequencies. Monkey N was trained on all three tasks and performed all three successfully as well (Figure 1C).

**Figure 1.**
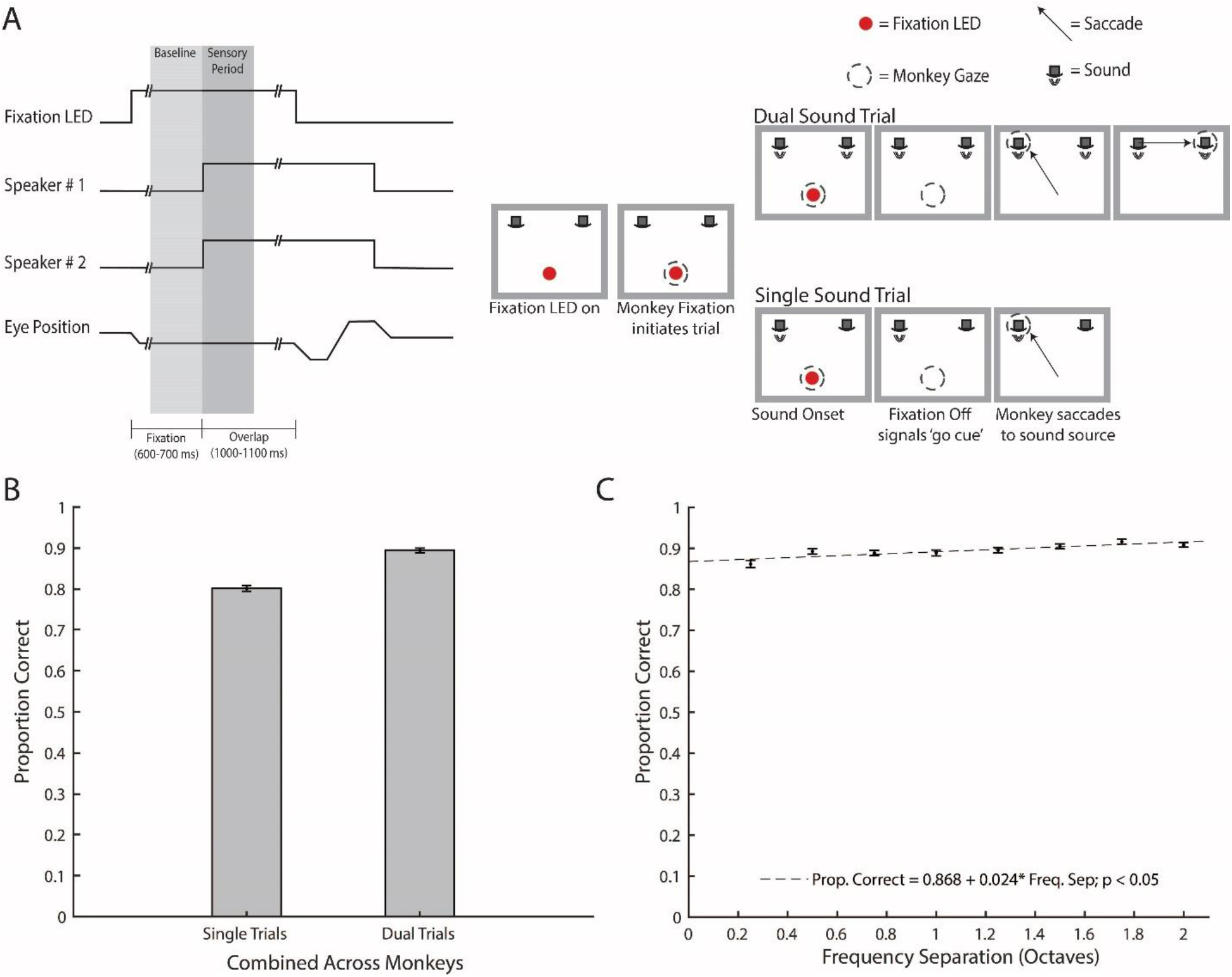
Task schematic and behavioral performance. (**A**) Monkeys were trained on a saccades-to-sounds task. Left panel: A time course schematic of the task. Speaker 2 onset would only occur on Dual sound trials and the second saccade and fixation would only be required if it was a Dual sound trial. Note, two sounds are targeted to one speaker on Control trials. Right Panel: In the saccades-to-sounds task the monkey would initiate a trial by fixating an LED. After variable interval fixation (600-700 ms), either 1, or 2 simultaneous, sound(s) were presented from one or two locations. After a variable interval of sound/fixation overlap (1000-1100 ms) the fixation light turned off, which cued the monkey to report to the origin(s) of the sound(s) via saccade(s). On Single sound and Control trials monkeys made a single saccade. On Dual sound trials monkeys made a sequence of saccades to the origin of one sound and then the other. (**B**) The combined proportion of correct Single sound trials (mean = 0.801 ± 0.007) and Dual sound trials (mean = 0.894 ± 0.0060) across both monkey Y and monkey N for 105 behavioral sessions collected during cell recordings. The control performance (mean= 0.721 ± 0.012) of monkey N across 60 behavioral sessions collected during cell recordings. (**C**) Change in performance across both monkeys as a function of the frequency separation between the two sounds for Dual sound trials. Performance, already near ceiling, nevertheless improves as distance between sound frequencies increases (significantly fit by a line with a slope > 0; slope 0.867, p <0.05). Error bars represent standard error of the mean (**B & C**).

#### Task details

During task performance, monkeys were seated in a dark, sound attenuated room with head movements restrained. Every session began with a series of visual trials in the four cardinal directions (up, down, left, and right) to permit calibration of the eye coil system. The speakers used for the current experiments were selected from a larger set ranging in position across +/− 24 degrees horizontally. This full set of speakers was used during initial animal training on saccade-to-sounds tasks involving single sounds; none of the speakers were visible to the monkeys during either training or the experiments conducted here.

The illumination of a fixation light (LED) at 0° in both the horizontal and vertical dimensions signified the start of the trial. Monkeys began the trial by acquiring and holding fixation on the light. After a variable duration of fixation (600-700 ms) either a single sound frequency or two simultaneous sounds of different frequencies would be presented while the monkey held fixation. After a variable overlap of fixation and sound presentation (1000-1100 ms), the fixation light was turned off, cueing the monkey to saccade to the location of the sound or sounds. The monkey had a brief time period for their eyes to leave the fixation window (700 ms) and enter the first target window (within 100 ms of leaving the fixation window). On Single sound and Control trials the monkey had to maintain fixation on this target for 300-650 ms. On Dual sound trials, the monkey had to hold fixation in the target window for at least 100 ms but complete a second saccade to the second target window within an additional 500 ms and maintain fixation in that window for 100-300 ms. The reinforcement window around the sound targets was 20 degrees horizontally, comparable to past studies of auditory-guided saccades (Metzger *et al.*, 2004).

Correct performance involved making saccades successfully to the appropriate number of targets within the appropriate time periods. This was almost impossible to do by accident, given the combination of spatial and temporal precision involved. If the monkeys performed the trial correctly by saccading into the correct windows at the correct times, they were rewarded with a grape juice solution (30% grape juice, 70% water). Reward was delivered through a drinking tube and controlled via a solenoid. Monkey Y (weight ∼ 3.7 kg) received approximately 0.1 milliliters of grape juice solution per trial and approximately 60 milliliters per session. Monkey N (weight ∼ 9.1 kg) received approximately 0.4 milliliters of grape juice solution per trial and approximately 400 milliliters per session. Both animals received additional fluid outside the sessions.

#### Stimulus conditions for neural recordings

In aggregate across the recording sessions there were 56 conditions, involving either one or two sound frequencies and locations. For simplicity, we will refer to the sound frequencies in two subsets, “Middle Frequencies” and “Flanker Frequencies”. The Middle Frequency sounds were used to assess the frequency response functions and consisted of the 8 middle sound frequencies (500 to 1682 Hz). The Flanker Frequency sounds flanked the Middle Frequency sounds, at either 420 or 2000 Hz. On Single sound trials any one of the 10 frequencies (8 Middle Frequency + 2 Flanker Frequency) could be presented in isolation at either of the ± 12° locations (n=20 conditions). On Dual sound trials, a Middle Frequency sound was presented at either ± 12° and paired with a Flanker Frequency sound at the opposite location. The Dual sound trials accounted for 32 conditions (8 Middle Frequency sounds × 2 paired Flanker Frequency sounds × 2 locations = 32). In addition to these 52 Single and Dual trial conditions, there were 4 Control trial conditions (1189 + either 420 or 2000 Hz, both frequencies at either ± 12°), which were presented to the animal for their behavioral importance but were not used further in the neurophysiological analysis.

For behavioral sessions prior to and between recording sessions monkeys were trained on all 52 (monkey Y – no Control trials) and 56 (monkey N) conditions with no restrictions on the spatial location of the Middle Frequency sounds. However, for individual recording sessions, we reduced the number of conditions to acquire enough trials per condition and to accommodate cell holding times (typically 1-2 hours). Specifically, we presented the Middle Frequency sounds at only one location, either ±12° but not both, on each day. The other stimulus conditions were unchanged: Flanker Frequency sounds were still presented at both locations on both Single and Dual stimulus trials, and Control stimulus conditions were presented at both locations as well. In net, each individual recording session consisted of 28 Single and Dual sound trial conditions: the Middle and Flanker Frequency sounds presented alone on one side (n=10), the Flanker Frequency sounds presented alone on the other side (n=2), the Dual sound trials involving pairing between the 8 Middle Frequencies on one side and either of the 2 Flanker frequencies at the other (n=8 × 2 = 16). In addition, the 4 Control conditions brought the total up to 32 conditions in any given session. The 28 Single and Dual conditions were used for the recording sessions involving monkey Y, whereas the Control trials were tested behaviorally only after the completion of the recordings in this animal. The full 32 Single, Dual, and Control conditions were used for the recording sessions involving monkey N. This counterbalanced design ensured a high degree of dissociation between locations and sound frequencies within and across days as well as an adequate number of trials for each tested condition: there were on average ∼24 attempted trials per tested condition for the recorded neurons in our dataset.

Conditions were randomly interleaved and the probabilities were weighted such that single sound and Dual sound trials occurred in a 1:1 ratio (45% each for monkey N, with the remaining 10% Control trials; 50% each for monkey Y). We took care to balance the number of stimuli presented at each location as follows: since the Dual stimuli trials each involved both locations, every frequency/location pairing was presented with equal likelihood. For Control trials, as noted above, both locations were used in all sessions and the frequency/location pairings could also be equally weighted. However, since on Single sound trials the Middle Frequencies within any given recording session were presented at only one location, we counterbalanced using the Flanker Frequencies. Flanker Frequencies were presented at both locations, with the number of trials at the side not used for the Middle Frequencies increased to make the likelihood of the +12° and −12° target locations about equal.

#### Training strategy

Monkeys underwent an initial training period in which they mastered a variety of visually-guided saccade tasks and became familiar with the laboratory setting and general nature of the experimental paradigm. Once acclimated to these tasks, tasks involving sounds were introduced. Making an eye movement to a sound is a natural behavior that typically occurs on the first trial provided the animal has not been previously habituated to sounds in the experimental rig (Metzger *et al.*, 2004). It remains to shape the animal to perform these eye movements consistently after the novelty has worn off. Monkeys were carefully reinforced for these critically important initial trials, and generally readily learned that sounds were as important as visual stimuli in the laboratory setting. Once the Single sound task was mastered, a variant of the Dual sound task in which the sounds were presented sequentially was introduced. Over time the inter-sound-interval was decreased until eventually the sounds were presented simultaneously (Caruso *et al.*, 2018). Monkey N was explicitly trained on all trial types; Monkey Y was only trained on the Single and Dual trial types but was tested on the Control trials after recording was complete (and performed single saccades on them).

#### Implementation

The saccades-to-sounds task used in this study was designed and implemented with proprietary software (Beethoven, RYKLIN Software Inc) and was adapted from a previous study (Caruso *et al.*, 2018). Eye position was acquired through an implanted scleral coil and Riverbend field system. Eye coil outputs were calibrated each day before behavioral and recording sessions. All behavioral data, including eye position data, were saved in Beethoven files for offline analysis.

### Neural Recordings

Neural activity was acquired through a Multichannel Acquisition Processor (MAP system, Plexon Inc). Single and multiunit activity signals were sent to an external speaker for auditory detection of neural activity. Once single units were isolated (using the box method, Plexon SortClient software), spike times were exported to Beethoven and saved for offline analysis.

A grid and turret system (Crist Instrument, 1mm spaced grid) was inserted into the chamber and used to hold the electrodes. Grid positions likely to intercept the IC were localized using structural MRI (Groh *et al.*, 2001). A single tungsten micro-electrode (FHC, ∼1-5 MΩ) was backloaded into a stainless-steel guide tube which was manually inserted approximately 1 cm into the brain and then advanced with a Microdrive (NAN Instruments) at 100-200 μm/s until the electrode tip was just outside of the guide tube. This was indicated by a mark on the electrode and confirmed by a change in the background activity typical upon electrode entrance into the brain. The depth on the microdive was then zeroed and the electrode was advanced at roughly 8 μm/s while monkeys sat passively listening to sounds of variable frequencies presented every few seconds. Once the expected depth and/or sound related activity was encountered in the background activity (as determined by ear via speaker and real-time plotting of multiunit response to sound onsets via Beethoven), electrode movement was stopped, and the electrode was allowed to settle for approximately 1 hour. The settling depth typically occurred 22 mm after exiting the guide tube. The electrode was then advanced in 1 μm increments until a cell was well isolated. If no cells were well isolated the electrode was retracted at roughly 8 μm/s.

A total of 105 neurons were recorded (monkey Y right IC, N=45; monkey N right IC, N=59; monkey N left IC, N=1). Forty-nine of the neurons had flanker frequencies presented at - 12° while the remaining 56 neurons had flanker frequencies presented at +12°.

### Analysis

#### Behavior

All behavioral analysis was done on attempted trials, defined as those in which the monkey successfully completed the fixation epoch and left the fixation window after the fixation light was turned off. Proportion correct was defined as the # of Correct Trials / # of Attempted Trials. Correct trials were defined as those in which the monkey made a saccade to the target window(s) of the played sound(s) (e.g. one saccade to either the ±12° on Single or Control sound trials or two saccades to ±12° on Dual sound trials). That is, on Single sound trials monkeys would make a single saccade to the appropriate speaker after the fixation LED turned off. On Dual sound trials monkeys could make a saccade to the left speaker then the right speaker or vice versa. Saccades to an intermediate location between the two sound locations on Dual trials (as might indicate the presence of summing localization; for review, see Blauert, 1983), were rare and classified as incorrect.

#### Neural Activity

All analysis of neural activity was done on correct trials and during the first 500 ms after sound onset, i.e. during a period of time when monkeys were fixating to reduce effects related to eye movements known to occur in the macaque IC (Figure 1A, “Sensory Period”) (Groh *et al.*, 2001; Porter *et al.*, 2006; 2007; Bulkin & Groh, 2012b; a). Neurons were classified as responsive if their spike count 500 ms after sound onset across all tested frequencies was significantly different from their spike count during baseline, the 500 ms prior to sound onset (t-test, p < 0.05). Ninety-three of the 105 neurons were classified as responsive. Neurons were classified as frequency selective, or tuned, if their spike counts 500 ms after sound onset were significantly modulated by sound frequency (10 Middle and Flanker sound frequencies from the same location, Single sound trials, one-way ANOVA, p < 0.05). Fifty-seven of 93 responsive neurons were classified as frequency selective. All cells that were included had peristimulus time histograms and frequency response functions that justified their inclusion by visual inspection.

#### Point Image

The point image refers to the proportion of recorded units that are responsive to a given condition (Capuano & McIlwain, 1981). Neurons were classified as responsive if, for a given condition, their spike count 500 ms after sound onset for that particular condition was significantly different than their spike count 500 ms prior to sound onset (t-test, p < 0.05).

#### d’ (d-prime)

To investigate how differently a cell responded to two sounds (here arbitrarily dubbed “A” and “B”), we calculated a sensitivity index or d’:

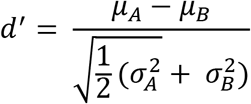

where μ_A_ and 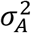 correspond to the mean spike count and spike count variance for single sound A, respectively, and μ_B_ and 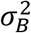 correspond to the mean spike count and spike count variance for single sound B, respectively. This metric quantifies the separation of the single sound A and single sound B spike count distributions in units of standard deviations. We took the magnitude of the d’ value (|d’|) since the order in the numerator is arbitrary (the A or B sound could be the first term). A |d’| value of 0 indicates the sound A and sound B spike count distributions are similar and higher values of |d’| indicate the sound A and sound B spike count distributions are more distinguishable.

#### Equivalent Rectangular Receptive Fields (ERRFs)

ERRFs serve as a measure of the sharpness of neural response functions. ERRFs were calculated in a similar fashion to previous work investigating spatial receptive fields (Lee & Middlebrooks, 2011), but applied here in the frequency domain. The area under the (non-baseline subtracted) frequency response function was measured with the *trapz()* function in MATLAB, which integrates the area under the frequency response function using the trapezoid method. This area was then divided by the peak firing rate observed across all frequencies, creating a measure of the width of the frequency receptive field normalized by peak firing rate.

#### Maximum-Likelihood Decoding

The decoding algorithm used here was adapted from previous work (Wallisch, 2014) and is similar to other implementations of maximum-likelihood decoding (Jazayeri & Movshon, 2006; Day & Delgutte, 2013). The goal of the decoder was to infer the stimulus that was presented on a set of held-out trials across all responsive units. For each run of the decoder, a held-out trial involving a particular condition was randomly selected for each unit (1 held-out trial of a given condition per cell, n = 93 cells, i.e. 93 held out trials per run). The mean spike counts for the remaining trials as a function of condition were then determined for each unit. The condition that was most likely to be associated with the spike count patterns on the set of held-out trials was then computed as detailed further below. The conditions to be decoded were: the 8 Middle Frequency sounds presented alone at one location (Single sound trials), the 2 Flanker Frequency sounds presented alone at the other location (Single sound trials), and the 16 Middle Frequency/Flanker Frequency combinations (Dual sound trials), for a total of 26 conditions. Note that the included data for Middle and Flanker Frequency sounds came from different locations across different neurons; these location differences were ignored in this analysis.

The predicted condition was the condition (1 out of the 26 possibilities) that had highest population-log-likelihood. This computation was repeated 1000 times per condition. Since there were not 1000 unique trials per condition for each cell, the particular held-out trial for a given neuron on a given repeat was sampled from all possible trials for that neuron with replacement. Because the neurons were recorded sequentially, variation in spike counts to a given stimulus condition is independent of the variation observed in other neurons, i.e. the variability in the responses of individual neurons to a particular stimulus condition on a particular day is not correlated with the responses of another individual neuron to that same stimulus condition on a different day. Spike counts were assumed to be Poisson-distributed (Caruso *et al.*, 2018).

To determine the population-log-likelihood, we first computed the likelihood of each condition at the cell level. Given a certain response on a held-out trial, the likelihood that a given condition elicited that response is computed from a Poisson distribution with an average and variance equal to the mean spike count (without the held-out data) for that condition. The likelihood for each of the 26 conditions given a certain spike count for a given cell is then given as:

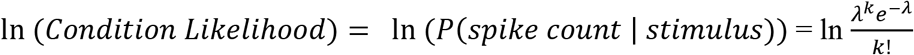

Where, λ is the mean spike count for the given condition and neuron and k is the held-out spike count. This results in a vector with 26 elements for each cell. Each element corresponds to the log-likelihood of that condition for that held-out spike count. To prevent occurrences of negative infinity, we changed instances of log(0) to log(0.1). These log-likelihood vectors were then summed across all cells to generate a population-log-likelihood:

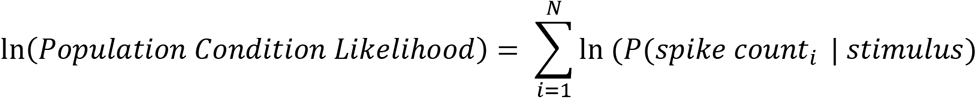

where *spike count* corresponds to the *ith* neuron spike count, and *N* is the number of neurons in the population. Therefore, for each repeat of the decoding computation, the result is a 26-element vector where each element corresponds to the population-log-likelihood for each condition. The predicted condition for that repeat is the element with the largest population-log-likelihood. Importantly, for any repeat the inferred condition could have been any one of the 26 conditions.

## Results

Our first step in evaluating the impact of sound combinations on the underlying neural code was to ascertain how well monkeys could distinguish the sounds perceptually. We found that monkeys could successfully report the locations of all sound frequency-location combinations used in this study. Across all attempted Dual sound trials, monkeys performed roughly 90% correct (Fig. 1B), compared to 80% correct on Single sound trials. This high degree of performance on the Dual sound trials was only modestly affected by the frequency separation between the two sounds: monkeys performed at about 87% correct for the smallest frequency separation of 0.25 octaves (Fig. 1C), rising slightly to 91% correct for the largest frequency separation of 2 octaves.

This good performance does not appear to be the result of recruiting largely unique sets of neurons to encode each of the two stimuli. Of the N=93 sound-responsive neurons we recorded (identified by t-test, see Methods), 45-70% responded significantly to any given stimulus presented individually (Fig. 2, red curve). The large proportion of responsive units suggests that sounds presented in this experiment recruit overlapping populations of neurons. The low frequency bias (negative slope of the red curve in Fig. 2) and large percentage of responsive units are similar to previous macaque IC findings (Bulkin & Groh, 2011).

**Figure 2.**
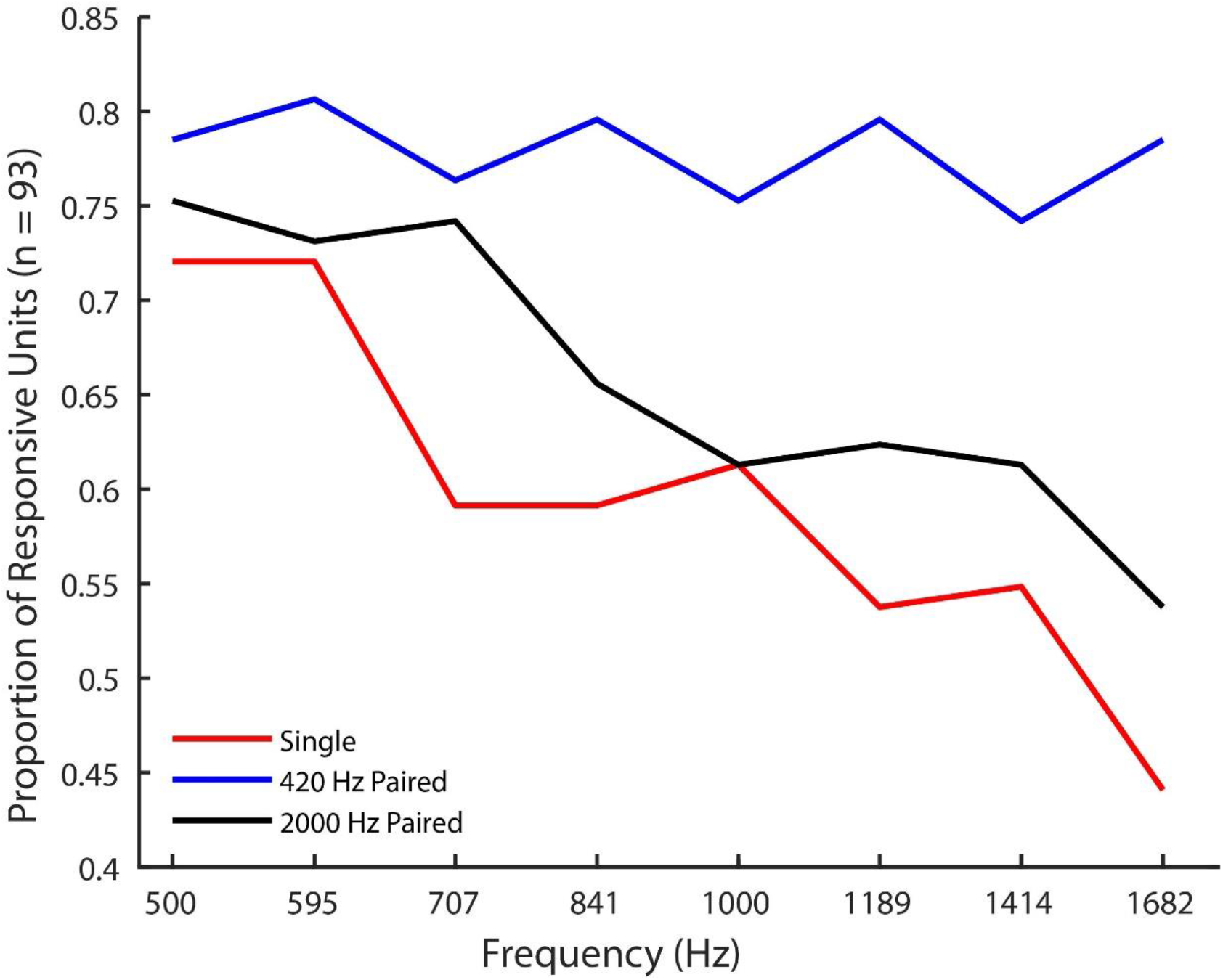
A point image plot, or the proportion of the responsive population that responds to a given frequency. Each point corresponds to the proportion of the responsive population that is significantly responsive compared to baseline for the given condition. A unit was counted as responsive if its firing rate in 0-500 ms after sound onset was significantly different than its firing rate from −500-0 ms before sound onset (t-test, p<0.05). Red corresponds to sounds presented in isolation, while blue and black correspond to Dual sound trials paired with 420 Hz or 2000 Hz, respectively. The x-axis corresponds to the frequency presented in isolation or paired with a particular Dual sound Flanker frequency.

When a second sound was added, more units responded (Fig. 2, blue and black curves vs. the red curve). This increase in the number of responsive units at the population level should not occur if individual units became more selective for sound frequency.

Although many neurons are responsive to a wide range of stimuli, it is possible that they still possess adequate selectivity to frequency to underlie this behavior, independent of any shifting or shrinking of their frequency response function. Therefore, we next compared the sensitivity of individual IC neurons to the component sounds of a given sound pair (arbitrarily called sound A vs. sound B). We quantified the separation of the sound A and sound B spike count distributions (e.g. a 500 Hz Middle Frequency vs. a 420 Hz Flanker Frequency) when either were presented in isolation) using a d’ analysis (see Methods).

Figure 3A shows the results for the comparison of Middle Frequency sounds with a 420 Hz Flanker Frequency sound. Units were sorted based on the sum of their |d’| values across conditions, with the units showing the highest aggregate |d’| on top. For the smallest frequency separation (left-most column), few neurons exhibit large d’ values: only about a third exhibit a |d’| value greater than 1 (bottom panel). Figure 3B shows the complementary pattern for the Middle Frequencies paired with the 2000 Hz Flanker Frequency sounds. Note that the unit orders in two panels are different; both are sorted so the unit with the highest summed |d’| within a panel is at the top. For both panels, a large minority of cells barely differentiated between any of the Middle and Flanker frequency sounds, as evidenced by the white bands at the bottom of the figure (roughly unit 70 to 93). We suspect the pattern in figure 3 is largely due to the majority of the frequency modulated neurons preferring low frequency sounds, as has been documented in several previous primate studies (Ryan & Miller, 1978; Bulkin & Groh, 2011). Overall, this analysis suggests that only a fraction of the recorded populations (roughly the top 20 units in both panels) possess sufficient sensitivity to underlie the improvement in performance due to frequency separation on Dual sound trials (Fig. 1C). Importantly, the results of figures 2 & 3 suggest that if successfully perceiving two sounds as distinct requires distinct populations of neurons to be active, frequency response functions would indeed likely need to shrink or shift to create such distinct peaks in a map for sound frequency.

**Figure 3.**
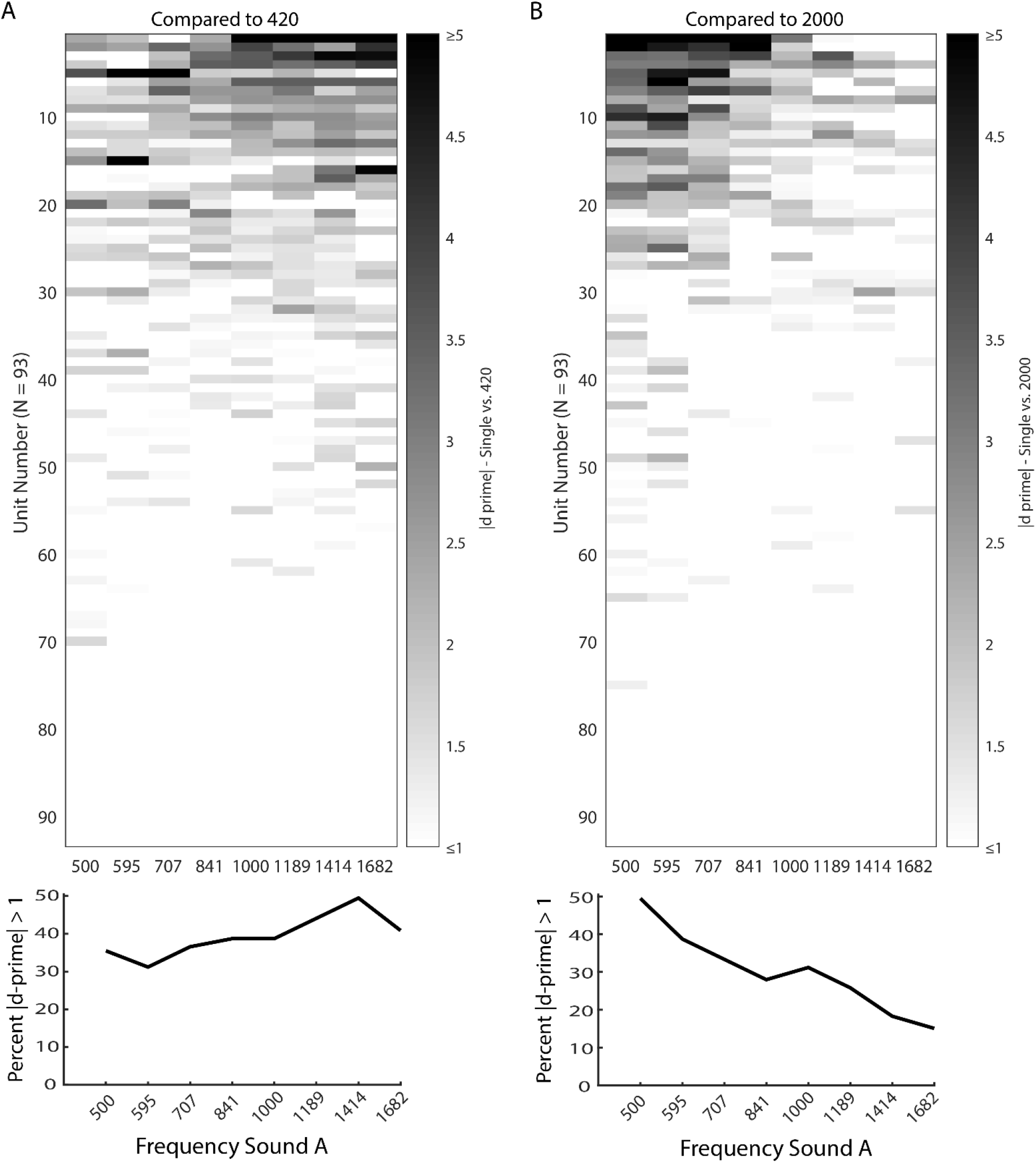
A sensitivity index, or |d’|, between two sounds of different frequencies when they are presented in isolation. (A) Top Panel: The magnitude of d’ between each of the 8 Middle Frequency “A” sounds and 420 Hz (“B” sound). A higher |d’| corresponds to a larger difference between the spike count distributions of the two sounds. Each row corresponds to a unit. Each column corresponds to a particular sound frequency. Units are sorted by their total |d’| across all 8 Middle Frequency sounds compared to 420 Hz. Bottom Panel: The percent of responsive neurons (N = 93) that have a |d’| value > 1 for each frequency compared to 420 Hz. (**B**) The same as panel (**A**) but compared to 2000 Hz. Note these units are sorted by their total |d’| across all 8 Middle Frequency sounds compared to 2000 Hz. Therefore, unit n in **A** does not necessarily correspond to unit n in panel **B**.

Therefore, we then evaluated whether neural response patterns change to facilitate the encoding of simultaneously presented sounds. Figure 4 compares the Single and Dual sound frequency response profiles of six example cells. On Single sound trials (red traces), neurons could prefer low frequencies (Fig. 4A, E, F), intermediate frequencies (Fig. 4B, D), or high frequencies (Fig. 4C). Concurrent sounds could change frequency response profiles in multiple ways. The neurons in figure 4A-C all become less responsive to sounds overall: the responses in the Dual sound cases were generally lower than the responses to the “better” of the two single sounds (e.g. black curves fall between the red and blue curves). Interestingly, the neurons shown in figure 4A & 4B are slightly inhibited by the Flanker frequency: the blue symbol and line are below zero. However, the decrement in firing rate on Dual sound trials when these Flankers are combined with Middle frequencies is a much larger effect. Some neurons appear to shift their preferred frequency (Fig. 4B & 4D), and some neurons appear unaffected by the presence of an additional sound (Fig. 4E, F). Interestingly, some neurons displayed multi-peaked receptive fields that were robust across single and dual sound conditions (Fig. 4F). Qualitatively, many neurons become less responsive in the presence of simultaneous sounds, but whether this suppression leads to frequency response profiles that sharpen or broaden requires quantification of response function changes across the population.

**Figure 4.**
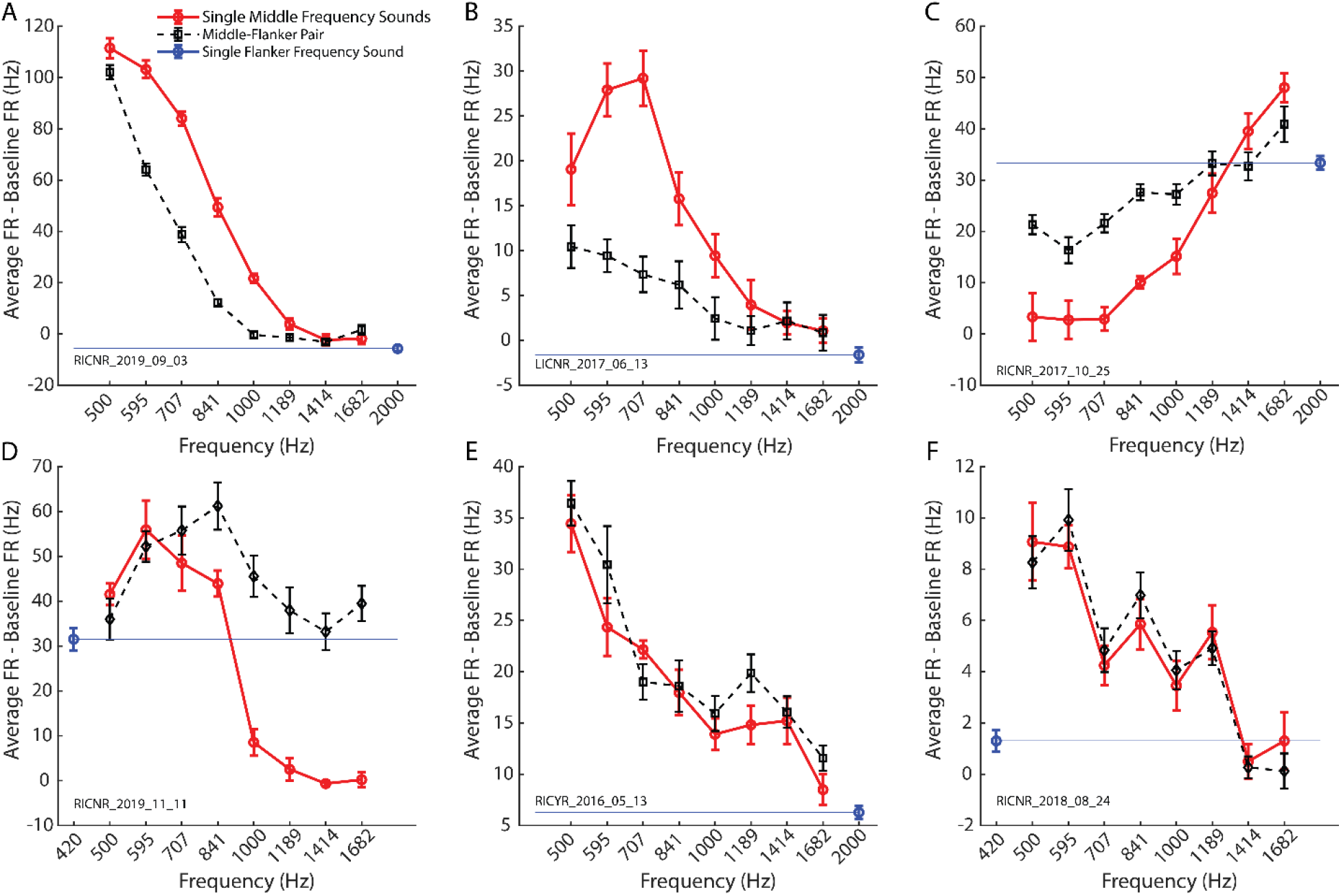
Example modulations of frequency response functions due to the presence of an additional sound in 6 different cells. (**A**) The response functions across conditions for a cell that displays sharpening in the dual sound condition. Each point corresponds to the baseline (−200 ms bin w.r.t sound onset) subtracted mean firing rate (500 ms bin w.r.t sound onset). Red points correspond to sounds presented in isolation. The blue point corresponds to the paired sound when presented in isolation. That is, the blue point is plotted at the flanker frequency used in the plotted Dual sound trials. The blue line is a reference line, so the firing rate of the paired sound is visible across the panel. The black points correspond to the dual sound condition. (**B and C**) The same as panel (**A**) but for two different cells that show a decrease in response and selectivity. (**D**) The same as panel (**A**) but for a cell that shifts its preferred frequency away from the paired sound. (**E & F**) The same as panel (**A**) but for two cells that seem to be unaffected by the presence of an additional sound. Error bars represent standard error of the mean (**A-F**). The series of letters and numbers in the bottom left of each panel is a unique session identifier for each neuron.

To quantify any neural response function changes across the population, we selected units that were selective to the Middle Frequencies (for example, the neurons shown in Figure 4B & 4D) when presented alone (N=57, one-way ANOVA, see Methods). The number of frequency-selective units dropped to N = 39 (∼68% of N = 57) when the Middle Frequency sounds were paired with the 420 Hz Flanker and N = 42 (∼73% of N = 57) when paired with the 2000 Hz Flanker. To evaluate effect size, we compared the F-statistic (the size of the effect of sound frequency on spike count) in the Dual vs. Single conditions (Figure 5A, B). If the neurons were more selective in the dual sound conditions, the F statistic should be higher for these conditions and the data would tend to lie above the line of slope one, but they do not. Instead, the data are shifted below the unity line, indicating that the bulk of the population tends to become significantly less frequency sensitive in both dual sound conditions (420 paired vs. single: Wilcoxon signed-rank test, p = 3.69 × 10^−6^; 2000 paired vs. single: Wilcoxon signed-rank test, p = 0.0253). To summarize, the frequency selectivity of the population of neurons was reduced by the presence of a concurrent sound, although few cells showed an enhanced selectivity (see points above unity line (Fig. 5A & B)).

**Figure 5.**
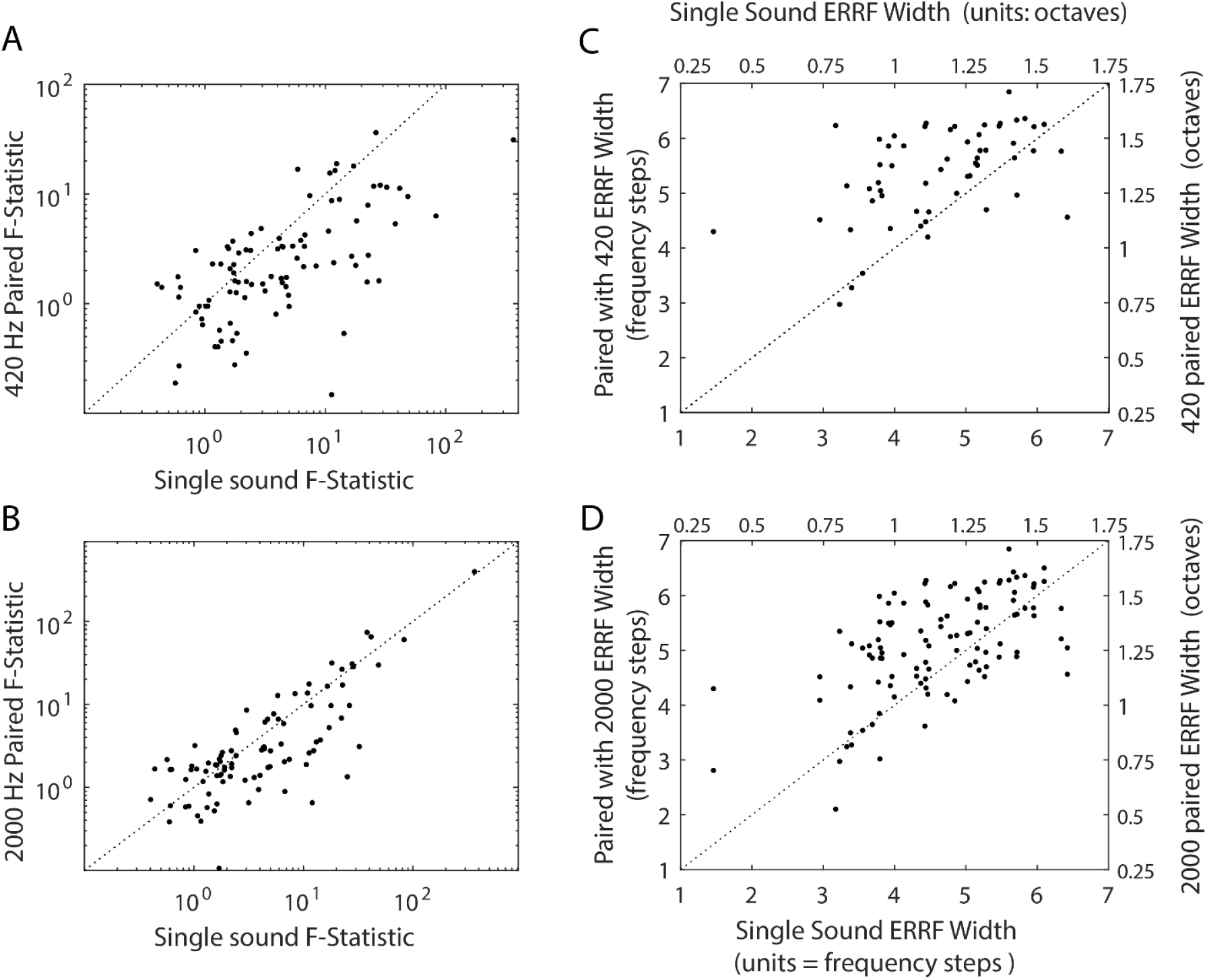
The population of neurons becomes less sensitive to frequency in dual sound conditions. (**A**) The F-statistic measured from a one-way ANOVA investigating firing rate across different frequencies (all 8 A sounds). A high F-statistic indicates that there are large differences in firing rate across the different frequencies. The single sound F-statistics are plotted against the 420 Hz paired F-statistics. Each point corresponds to one unit, out of 57 frequency selective units. Points above the unity line indicate that frequency modulates firing rate more in the 420 Hz paired condition. Points on the unity line indicate there is no change in the modulation of firing rate across the two conditions. Points below the unity line indicate that frequency modulates firing more in the single sound condition. (**B**) The same as (**A**) but for compared to the 2000 Hz paired condition. (**C**) The width of the ERRF for each unit in the single sound condition plotted against the width of the ERRF for each unit plotted in the 420 Hz paired condition. Each point represents a unit, out of 57 frequency modulated cells. The dashed line indicates unity, or ERRFs of equal width in the single sound and 420 Hz paired condition. Points above unity indicate ERRFs of greater width in the dual-sound condition. Points below unity indicate ERRFs of greater width in the single-sound condition. (**D**) The same as (**C**) except for the 2000 Hz paired condition.

The preceding analysis considers how the means and variances in firing depend on sound frequency but does not directly investigate whether frequency response functions change in systematic ways, such as changes in the width or peak (best frequency). To investigate possible changes in frequency response function width, we implemented the equivalent rectangular receptive field analysis (ERRF) from Lee and Middlebrooks (2011); see also Burton *et al.* (2018). Briefly, the ERRF divides the area under the response function by the peak firing rate, creating a metric that incorporates the number of frequencies to which the unit responds based in part on the level of activity evoked by each of those stimuli. For example, many of the cells display a suppression in the dual sound condition (Fig. 4 A-C), yet these suppressions seem to be of different kinds. Generally, suppressions that affect firing rate across all conditions (Fig. 4B & C) tend to increase ERRF width while suppressions that affect some conditions but leave the peak of the measured response untouched (Fig. 4A) tend to decrease the ERRF width.

To compare frequency response function width between single and dual sound conditions, the distributions of each were plotted against each other for all frequency selective neurons (N = 57, Fig. 5C & D). If the ERRF widths were similar across single and dual sound conditions the distribution would fall upon the unity line (black line in Fig. 5C & D). If the ERRF widths were narrower in the dual sound condition the distribution would be shifted below unity, and if the ERRF widths were narrower in the single sound condition the distribution would be shifted above unity. It is quite apparent that ERRF widths are significantly broader in the dual sound conditions, for both high- (Wilcoxon signed-rank test, p = 0.021) and low-frequency paired sounds (Wilcoxon signed-rank test, p = 1.82 × 10^−7^). In summary, across the population the majority of neurons actually broaden their response functions in response to presentation of two simultaneous sounds.

Could some neurons shift their frequency response function, exhibiting a different best frequency in response to dual sounds (e.g. Fig 4D)? To investigate this possibility we selected neurons with firing rates significantly modulated by frequency and with single sound best frequencies that fell within the middle six A sounds (595-1414 Hz), leaving out the edge cases for which a shift outside the tested range could not in principle be observed (N=17). We then compared their single sound best frequency to their best frequency in the dual sound conditions. Seventeen neurons met these criteria. Figure 6 shows how best frequency changes in the 420 Hz paired condition (Fig. 6A) or the 2000 Hz paired condition (Fig. 6B). Even though some individual neurons appeared to shift their best frequency, we did not observe a consistent overall mean shift in the population. The slight shift of the mean best frequency away from the paired sound was not significant for either the 420 Hz dual sound (single mean = 759 ± 76 Hz S.E.M. vs. dual mean = 938 ± 114 Hz S.E.M., paired t-test, p = 0.072) or the 2000 Hz dual sound conditions (2000 dual mean = 741 ± 76 Hz S.E.M., paired t-test, p = 0.56). These findings suggest that shifting of response functions is not a ubiquitous computation within the population and is unlikely to substantially contribute to the encoding of simultaneous sounds.

**Figure 6.**
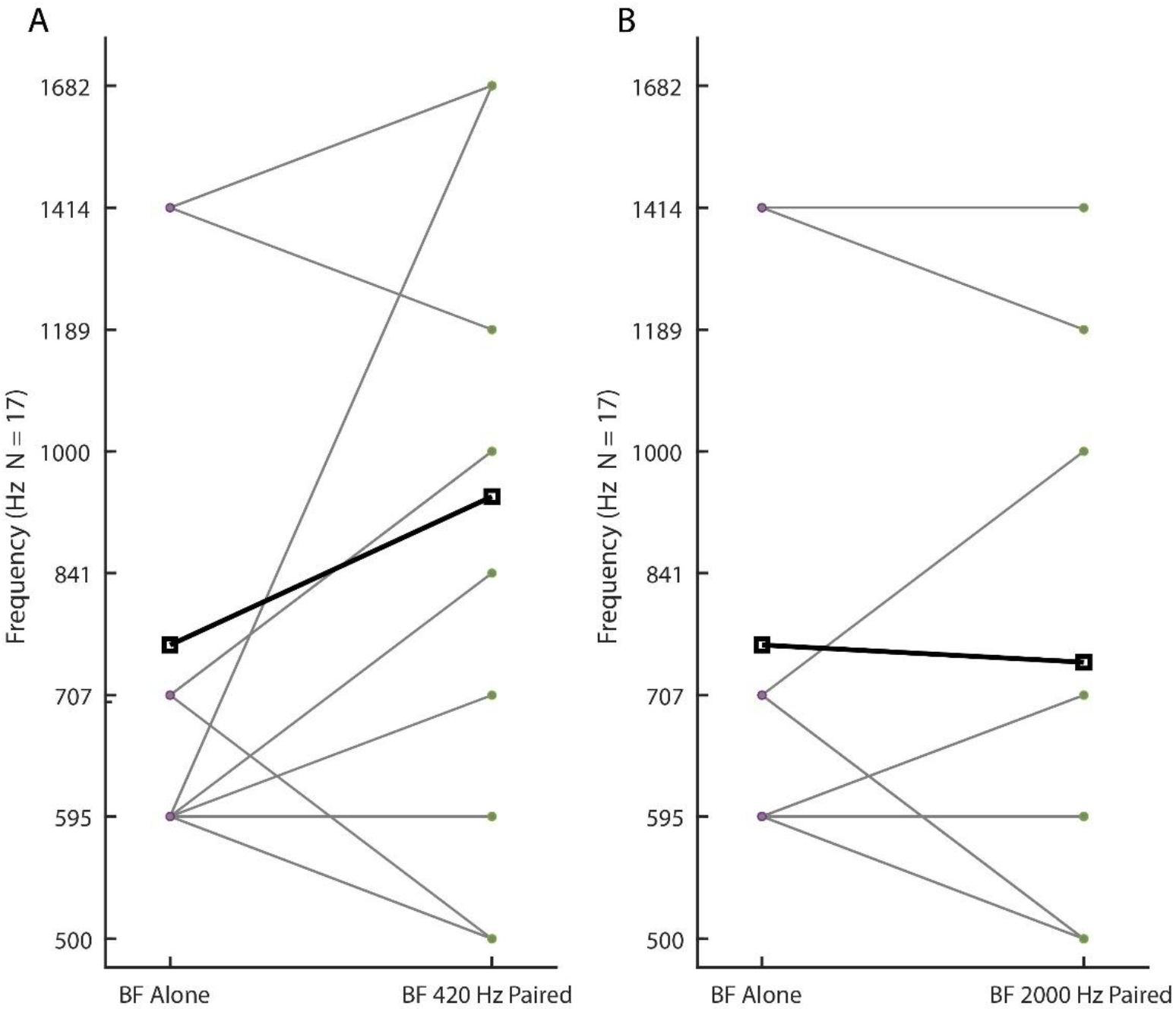
The presence of an additional sound does not cause a large shift in best frequency (BF) across the population of recorded cells. (**A**) The single sound best frequency plotted against the 420 Hz paired best frequency in the 17 cells that have best frequencies between 595-1414 Hz (grey lines). The bold line indicates the mean best frequency across the population. There is no significant shift in best frequency across the two conditions (single mean = 759.3 ± 76.48 S.E.M. vs. dual mean = 937.6 ± 113.6 S.E.M., paired t-test, p = 0.072). (**B**) The same as (**A**) but for the 2000 Hz paired condition. There is no significant shift in best frequency across the two conditions (single mean = 759.3 ± 76.48 S.E.M. vs. dual mean = 740.9 ± 75.67 S.E.M., paired t-test, p = 0.56).

In short, the modulations of frequency response functions on Dual sound trials do not appear to be optimized for the preservation of information about either sound. However, it remains to be seen if these cell-by-cell effects substantially affect the amount of frequency information in the population as a whole. To determine the impact of multiple sounds on the efficacy of coding stimulus related information across the population of recorded IC neurons, we evaluated how this code might be read out.

We first used a population-maximum-likelihood decoder to infer the condition from the observed spike count on a set of held-out trials (see Methods). Spike counts were assumed to be Poisson-distributed (Caruso *et al.*, 2018), and neurons were assumed to be conditionally independent of each other (a reasonable assumption given they were recorded at different times). For each held-out trial, the log-likelihood for each condition was summed across the population and the condition with the maximum-population-log-likelihood was predicted to be the held-out trial’s condition. This computation was repeated 1000 times per condition.

Overall, the decoder inferred every condition above chance (16 dual sound conditions, 10 single sound conditions for a total of 26 conditions, chance = 1/26, see methods). Figures 7A, C, and E show the proportion of correctly predicted conditions across the 3 paired sound trial-types, sounds presented in isolation, sounds paired with 420 Hz, or sounds paired with 2000 Hz, respectively. Qualitatively, the mean across all sounds presented in isolation (∼66.7% correct) was higher than the mean across all 420 Hz paired conditions (∼47.1% correct) or the mean across all 2000 Hz paired conditions (∼51.8% correct). Figures 7B, D, F display the average population likelihood functions predicting the held-out single sound, 420-paired, or 2000-paired trials, respectively. Each held-out condition has one likelihood function (e.g. held-out 420 trials correspond to the solid black line in Fig. 7B) that shows how likely each of the 26 conditions is for that held-out data. Therefore, if the population, on average, correctly predicts the frequency presented on the held-out trials, the log-likelihood should be the highest for that condition (e.g. the maximum of the 420 Hz likelihood function occurs at 420 Hz). The likelihood functions show that the confusion in decoding tends to come from neighboring frequencies (Fig. 7B, D, F). For example, for trials with a held-out condition of 841 Hz (Fig. 7B), the highest likelihood is for 841 Hz (solid grey line) and the next two most likely conditions are 707 Hz (black dashed line) and 1000 Hz (grey dashed line). This can be further seen in the confusion matrix (Fig. 8A) where the off diagonals are slightly darker in the dual compared to single sound conditions. This confusion arises due to the increase in likelihood when a low frequency sound is present (420 Hz paired and when 2000 Hz is paired with a low frequency: Fig. 8B), which may stem from the population’s low frequency bias (Fig. 2; Bulkin & Groh 2011). Ultimately, the changes in the frequency response functions seem to degrade the amount of information available in the population concerning each sound, resulting in worse decoding on Dual sound trials.

**Figure 7.**
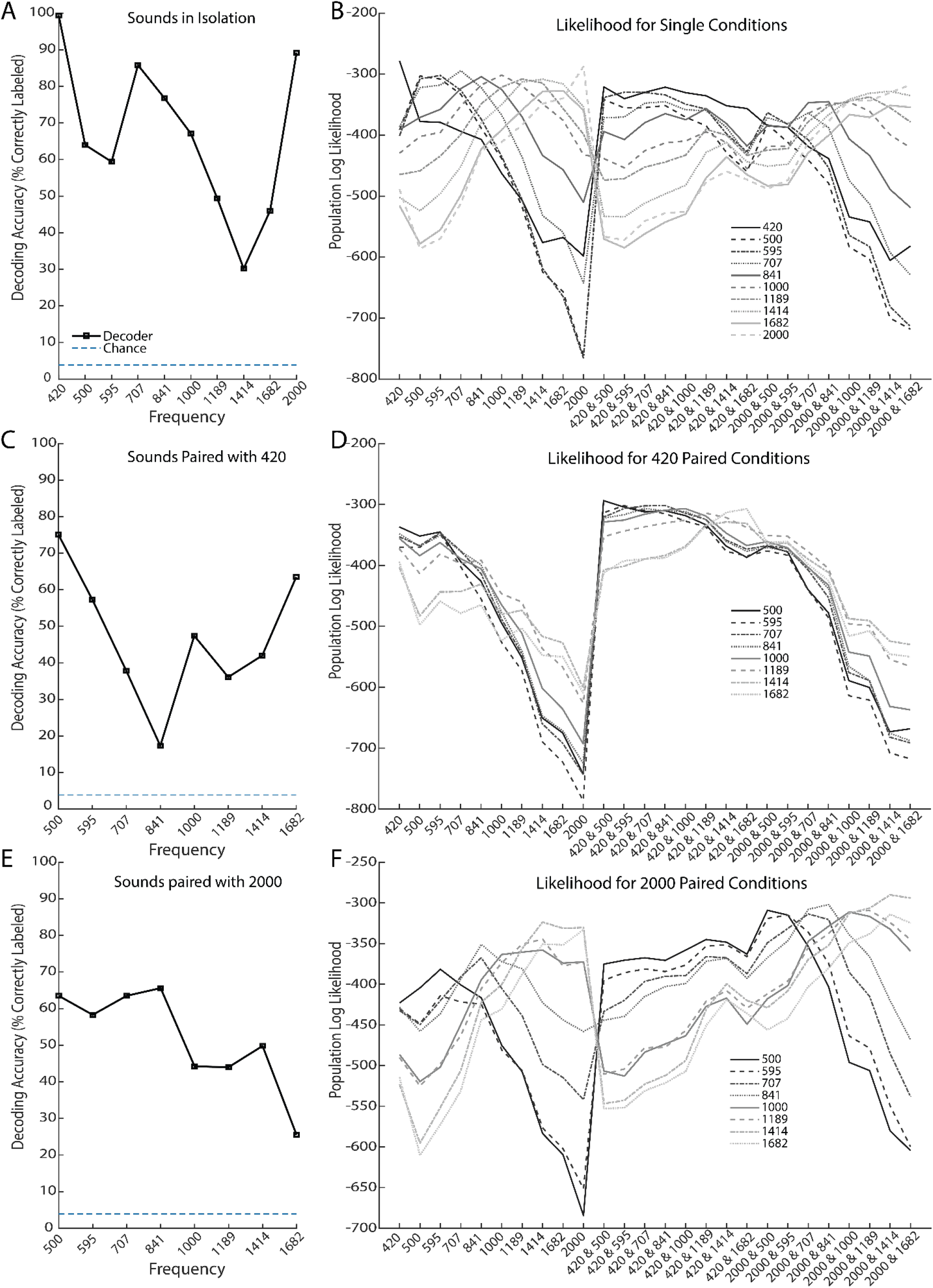
Accuracy of population maximum-log-likelihood decoding and mean log-likelihood functions across conditions. (**A**) The accuracy of the population (N = 93) maximum-log-likelihood decoder across each single sound condition. Each point corresponds to the percent of correctly labeled repeats (out of 1000) for each condition. Chance level prediction is 1/26 or ∼ 3.8% (**B**) The mean population log-likelihood function for each condition across all potential predicted conditions. Each condition had the chance of being labeled as any one of the 26 single or dual conditions. (**C**) The same as panel (**A**) but for 420 Hz paired conditions. (**D**) The same as panel (**B**) but for 420 Hz paired conditions. (**E**) The same as panel (**A**) but for 2000 Hz paired conditions. (**F**) The same as panel (**B**) but for 2000 Hz paired conditions.

**Figure 8.**
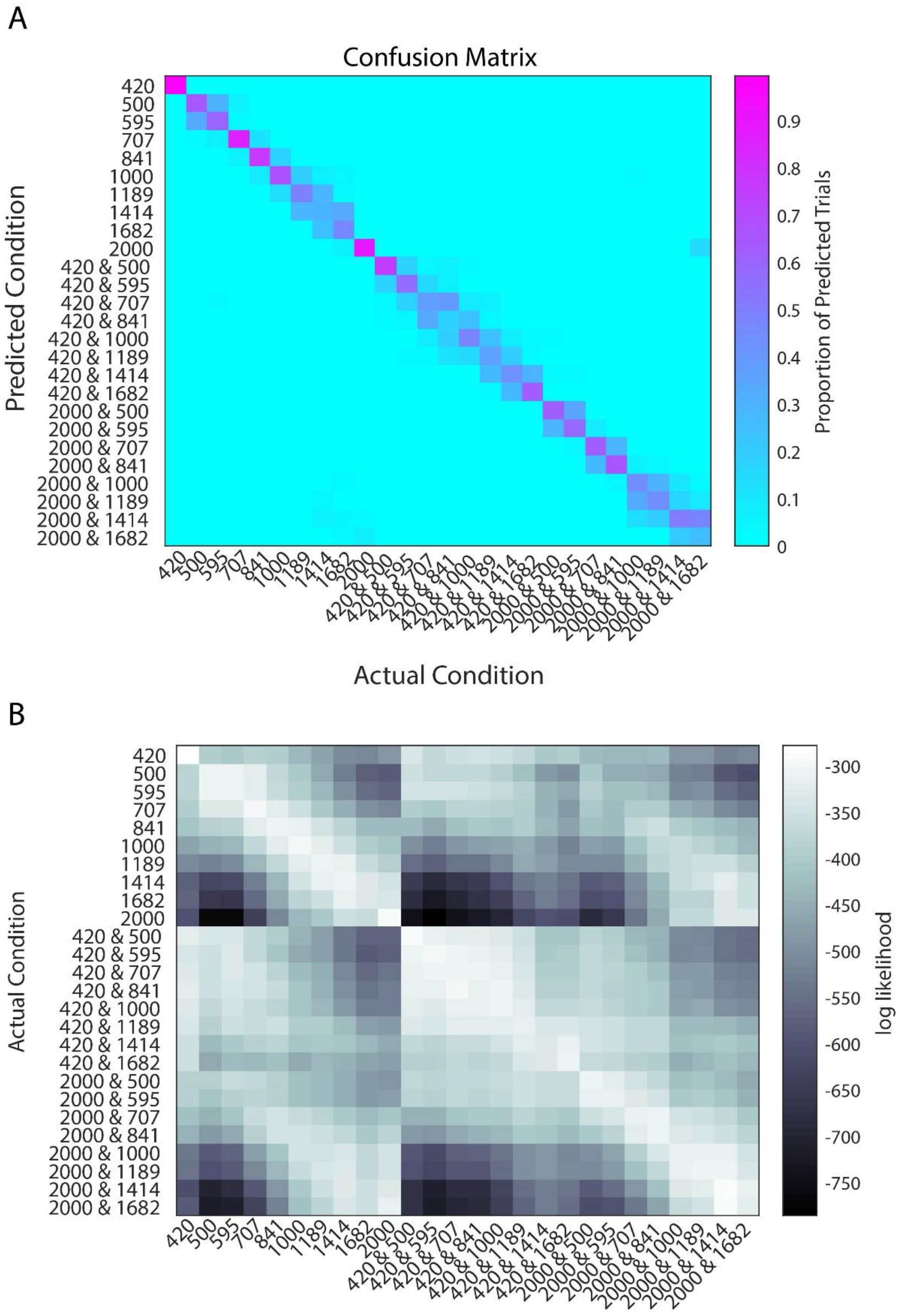
Confusion and mean-likelihood matrices for the decoder. (**A**) The confusion matrix for the decoder. Each column corresponds to the actual condition and each row corresponds to the predicted condition. The color of each square is the proportion of predicted trials (out of 1000) that fell into that bin. The diagonal corresponds to correct predictions. (**B**) A matrix of mean log-likelihood across conditions for the decoder. Each row corresponds to the likelihood function for that condition while each column corresponds to the mean log-likelihood for that condition across all actual conditions (e.g. leftmost column corresponds to the mean log-likelihood for 420 Hz across all conditions).

Until now, all the analyses presented probed how the encoding of sound changes in the presence of an additional sound. With the implementation of the maximum-likelihood decoder, we can now ask if single and dual sounds are encoded with similar neural codes, as is typically assumed in experiments only presenting a single stimulus. To investigate if the monkey IC uses a similar code for the single and dual sound conditions, we attempted to decode the dual sound condition using the mean responses observed on the Single sound trials. Each held-out Dual sound trial for a given Flanker Frequency could, therefore, be labeled as one of the 8 Middle Frequency sounds. Performance was markedly worse in the 420 Hz condition when decoded with the single sound response patterns (accuracy = 16.5%; chance 12.5%) compared to the previous decoder that had access to the dual sound response patterns (accuracy ∼ 47.1%; chance ∼ 3.8%). This is evident in the confusion matrix (Fig. 9A) which shows 595 Hz is the most predicted stimulus for seven of the eight conditions. This is likely due to the low frequency preference of the monkey IC (Bulkin & Groh 2011). The performance in the 2000 Hz condition when decoded with the single sound response patterns (accuracy = 40.65%; chance = 12.5%) is also worse compared to the previous decoder that had access to the dual sound response patterns (accuracy ∼ 51.8% correct; chance ∼3.8%). Unlike the 420 Hz paired condition, the 2000 Hz paired conditions seem to confuse neighboring frequencies (Fig. 9B). These results show that the IC would need to use different coding strategies in the single and dual sound conditions to maintain information about sound frequency, implying an IC readout may need to implement a different set of weights depending on the number of sounds present.

**Figure 9.**
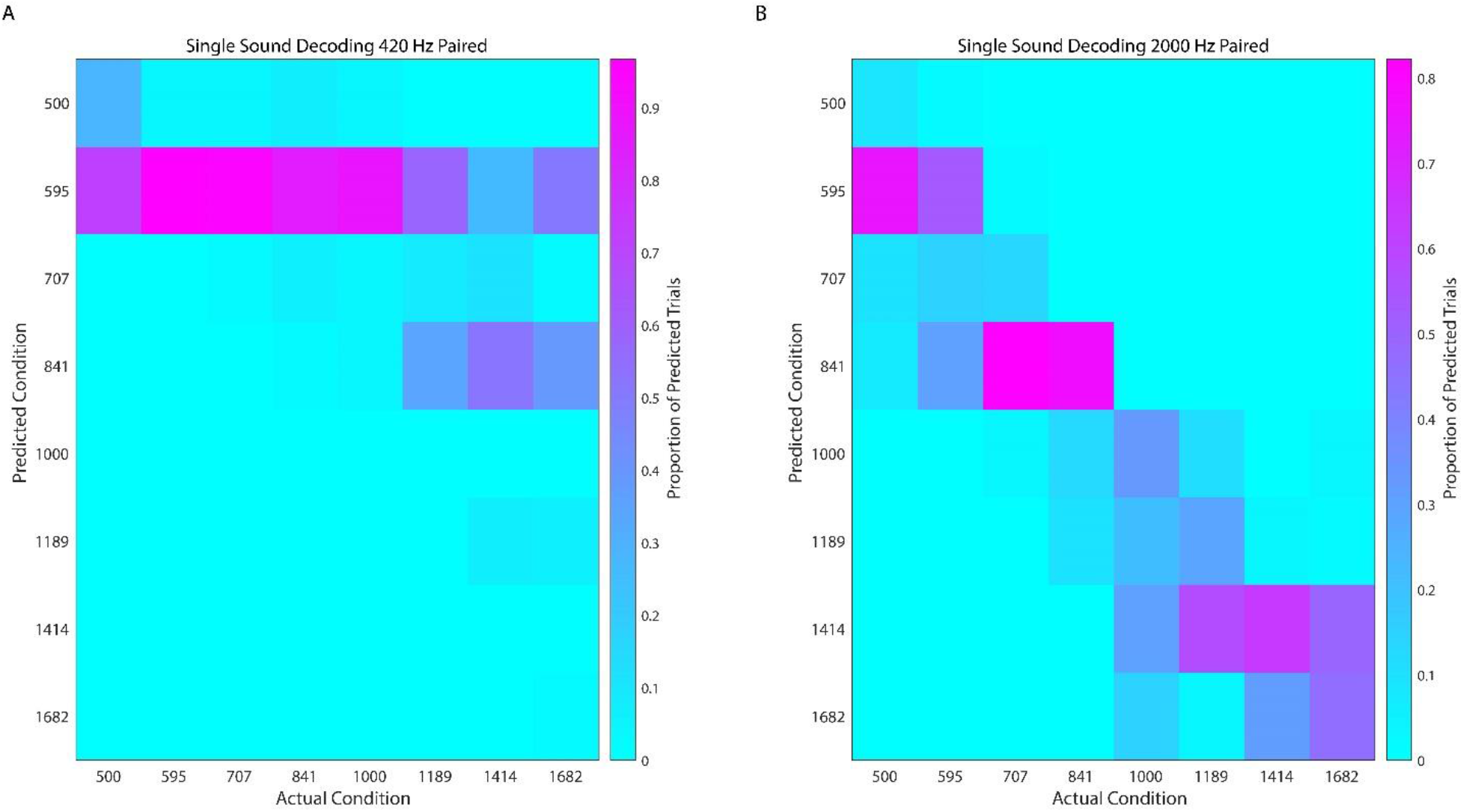
Decoding the paired condition with the single sound response patterns impairs performance. (**A**) The confusion matrix for decoding 420 Hz paired conditions using the response patterns observed on single sound trials. Each column corresponds to the actual condition and each row corresponds to the predictable condition. The color of each square is the proportion of predicted trials (out of 1000) that fell into that bin. The diagonal corresponds to correct predictions. (**B**) The same as (**A**) but for decoding the 2000 Hz paired conditions.

Finally, we considered whether the inferior colliculus population activity contains information about the number of sounds present, regardless of their frequency spectra? To answer this question, we used our maximum-likelihood decoder to not decode frequency, but rather to decode out the number of sounds on a given trial. The decoder was quite successful at decoding the number of sounds, accurately classifying 91.8% of Single sound trials and 94.9% of Dual sound trials -- much higher than the chancel level of 50%. This is somewhat unsurprising given the presence of an additional sound affects the firing rate of many cells in a consistent fashion (for example, Fig. 4A-D). Thus, although the modulations to individual neural response functions lead to the IC population to lose information about sound identity, a representation of the number of sounds occurring on a given trial appears to be preserved.

## Discussion

How the brain simultaneously encodes multiple items, particularly when more than one item falls within a given neuron’s receptive field, is an often-overlooked problem in neuroscience. Here we tested one possible neural solution to this problem: that response functions might change in ways that reduce ambiguity and preserve information about both sounds at the population level. We first verified that monkeys could successfully make saccades to each of two concurrently presented sounds whose center frequencies were separated by as little as 0.25 octaves (and whose frequency content overlaps). We then investigated how frequency response functions were affected by the presence of that second sound. We found that the sounds we presented recruited largely overlapping populations (Fig. 2 & 3), and that on Dual sound trials the majority of these cells’ firing rates were less modulated by sound frequency and had their response functions broadened compared to those observed on Single-sound trials (Fig. 5). There was little evidence of systematic shifts in best frequency (Fig. 6).

Overall, these changes in frequency response functions reduce the information available to the maximum-likelihood decoder, decreasing the decoding accuracy in the dual sound conditions (Fig. 7 & 8). The poor decoding performance when using the single sound response patterns to decode dual sound conditions (Fig. 9) suggests that a different read-out is needed depending on the number of sounds – a startling possibility given that the brain cannot have prior knowledge of how many sounds are in the environment except by virtue of what it detects via sensory input. Conceivably, the brain could use a two-step decoding process, one to assess the number of sounds (which our maximum likelihood decoder was successful at), and a second to deploy a sound-number-specific readout based on the results of the first step. However, such an operation is cumbersome to envision even for two sounds, and scaling up to a larger number seems implausible.

More broadly, a quantitative accounting for perceptual abilities in light of the coarseness of coding is an important unsolved problem in sensory systems. In the auditory system, many reports highlight the presence of some neurons with “sharp” tuning, especially to sound frequency (e.g. Sadagopan & Wang, 2010). It is possible that a very small number of very sharply tuned neurons are responsible for the monkeys’ perceptual abilities. However, there is a discrepancy between what “sharp” means neurophysiologically and what is observed psychophysically. For example, the median bandwidth of marmoset auditory cortical neurons ranges from 0.25-.5 octaves (Sadagopan & Wang, 2008); cat and rhesus monkey appear broadly comparable (Calford *et al.*, 1983; Recanzone *et al.*, 2000) . These values are an order of magnitude higher than the 0.03 octave frequency discrimination ability in the 2 kHz range that has been reported for monkeys perceptually (Sinnott *et al.*, 1987). It is also worth noting that both the perceptual and neurophysiological measures of frequency sensitivity mentioned above involved sounds in isolation; representations of frequency are known to be affected by the presence of noise (Miller *et al.*, 1987).

It is not always clear how to relate such metrics regarding individual neurons to the population level and thence to perception. The bandwidth metrics used in most neurophysiological studies are individualized for each neuron’s sound threshold level (e.g. Q10, which refers to the bandwidth of a frequency tuning curve 10 dB above threshold for that neuron, (Kiang *et al.*, 1965). A further difficulty is that definitions of “threshold” for a neuron may vary depending on the analysis methods, making it hard to generalize across studies. More recent reverse correlations methods using broad band sounds represent a useful advance in determining robust quantitative measures of frequency tuning (Versnel *et al.*, 2009; Eggermont, 2011; David, 2018). Overall, population-level metrics that describe neural response patterns for a fixed stimulus set tested in every neuron are needed to advance efforts to account for perception at the neural level.

To our knowledge, this is the first computational analysis of how sound frequency information is decoded from the responses of neurons in the inferior colliculus of awake monkeys performing a behavioral task. Our findings confirm that the coarseness of coding is indeed a problem: even in the single sound condition, our decoding analysis fell well short of perfect performance, and performed at roughly 30% for the worst condition (frequency 1414, Fig. 7A), far worse than the monkeys’ actual performance of ∼90% correct. The decoding analysis also highlights a general limitation of previous decoding studies, namely that performance assessed under single-stimulus conditions does not necessarily generalize well to the more natural multi-stimulus case.

Our results seem to be in-line with prior work showing degradation of information when there are multiple stimuli presented (Day *et al.*, 2012; Day & Delgutte, 2013; Henry & Kohn, 2020). If this reduction in information scales with number of stimuli it could potentially explain the finding that the number of distinctly identifiable sounds saturates at three (Zhong & Yost, 2017).

Importantly, these results refute the primary hypothesis we set out to test: the notion that the changes in frequency response functions could be used to overcome the multiplicity problem, and suggest that alternative coding possibilities should be explored. In particular, we recently proposed a novel theory of neural representations in which response functions could stay unchanged during presentation of multiple stimuli, but neurons might instead alternate between encoding one stimulus and the other, allowing both stimuli to be represented successfully across time (Caruso *et al.*, 2018; Glynn *et al.*, 2020; Mohl *et al.*, 2020) . We developed novel statistical methods to test this possibility, and found evidence in support of it in both IC neurons and neurons in a visual cortical area. Specifically, Caruso et. al. (2018) found monkey IC neurons that respond to combinations of sounds ‘A’ and ‘B’ as if only sound ‘A’ was presented on some trials and only sound ‘B’ on other trials. Additionally, there were also some cells that alternated their firing rates between that of sound ‘A’ and sound ‘B’ within a given trial, potentially allowing downstream neurons to represent each sound across time and across the neural population (Caruso *et al.*, 2018).

Such activity fluctuations may well have occurred in the present study. To better compare with previous literature, we made the simplifying assumption here that the time-and-trial-pooled average response of a neuron to a combination of stimuli is reflective of the information that the neuron encodes. If the underlying activity on Dual sound trials actually fluctuates between the A-like and B-like response patterns (as seen in Caruso *et al.*, 2018), the time-and-trial-pooled average will be a poor measure of the information present in neural signals. The fact that average responses to AB sounds were generally between the average responses to A and B sounds presented alone supports the possibility that fluctuations may underlie at least a portion of the results observed here. Future work will test this possibility.

In principle, changes in frequency response functions as investigated here and fluctuating activity patterns as investigated previously (Caruso *et al.*, 2018) both have the potential to limit the degree to which a given neuron is faced with the task of encoding more than one stimulus at a time. It is also possible that alternative forms of coding such as patterns of spike timing or first spike latency contribute (e.g. Furukawa & Middlebrooks, 2002; Chase & Young, 2006); these were not explored in our study which focused on total spike count in a broad temporal epoch, 500 ms, a window that encompassed both transient and sustained aspects of sound-evoked activity (Ryan & Miller, 1977; Bulkin & Groh, 2011). It is therefore conceivable that better decoding accuracy might occur when using a different temporal epoch or metric of neural activity.

It also remains possible that information-preserving changes in frequency responses might be more evident when tested with a wider range of sound frequencies, allowing a fuller exploration of frequency response functions. Our small number of frequencies could potentially explain why many of our neurons appear to have monotonic frequency response functions. We suspect with more frequencies these receptive fields would be circumscribed and would display proper broadening; currently we use the term broaden to describe monotonic frequency response functions that flatten, or have less change in firing rate per change in sound frequency. However, the granularity of our testing, 0.25 octave spacing, was clearly coarser than monkeys’ perceptual abilities, and the overall range, 2 octaves, should have been adequate to demonstrate an effect if changes in frequency tuning were a major contributor to these perceptual abilities.

In conclusion, our study reveals important shortcomings in neural representations of multiple sounds in the primate IC. Outside the rarefied environment of the laboratory, more than one sound is the rule, not the exception. Animals, including humans, are capable of perceiving such sounds as distinct from one another, suggesting that new forms of neural coding should be explored to account for these perceptual abilities.

## Abbreviations

ERRFs: Equivalent Rectangular Receptive Fields
IC: Inferior Colliculus

## Acknowledgements

We are grateful to Stephen Lisberger, Lindsey Glickfeld, Henry Greenside, Patrick Mayo, Christopher Henry, Jeff Mohl, Surya Tokdar, John Pearson, Stephanie Schlebusch, Meredith Schmehl, Gelana Tostaeva, Justine Griego and members of the Groh lab for comments on this work. We are also grateful to Valeria Caruso for her help in establishing the behavior paradigm. This work was funded through NIH grants DC016363 and DC013906 to JMG.

## Author Contributions

SMW and JMG designed the experiments. SMW collected and analyzed data and wrote the manuscript. SMW and JMG edited the manuscript. JMG acquired funding for this project.

## Data Availability Statement

Data can be made available upon reasonable request.

